# Covalently linked peptides and membrane potential enable CyaA segment translocation

**DOI:** 10.64898/2026.04.22.716334

**Authors:** Gaia Scilironi, Nicolas Carvalho, Jacinthe Frangieh, Corentin Léger, Dorothée Raoux-Barbot, J. Iñaki Guijarro, Daniel Ladant, Sophie Cribier, Nicolas Rodriguez, Alexandre Chenal

## Abstract

The adenylate cyclase toxin (CyaA) from *Bordetella pertussis* intoxicates host cells by directly translocating its N-terminal catalytic domain across the plasma membrane; however, the forces driving this unique process remain poorly defined. Here, we dissect the membrane translocation mechanisms of two peptide segments derived from CyaA: P233 and P454 from the catalytic domain and the translocation region, respectively. Both P454 and P233 are calmodulin-binding segments that are sequentially involved in the translocation and activation of the catalytic domain. Using a newly developed Droplet Interface Bilayer (DIB) approach, called DIB-Pipette, which enables direct visualization of peptide transport under controlled membrane potentials, we show that P454 translocates across membranes independently of membrane potential, whereas P233 translocation requires a negative electric membrane potential. Strikingly, covalent coupling of P233 and P454 enables efficient translocation of the resulting peptide even in the absence of a membrane potential. Together, these results suggest that two distinct membrane-active segments within CyaA act cooperatively to promote translocation at the peptide level, revealing an intrinsic mechanism that may contribute to membrane potential-dependent translocation. These findings provide new mechanistic insights into CyaA cell intoxication process and reveal a multifunctional strategy for protein delivery across membranes.

**Graphical abstract:** 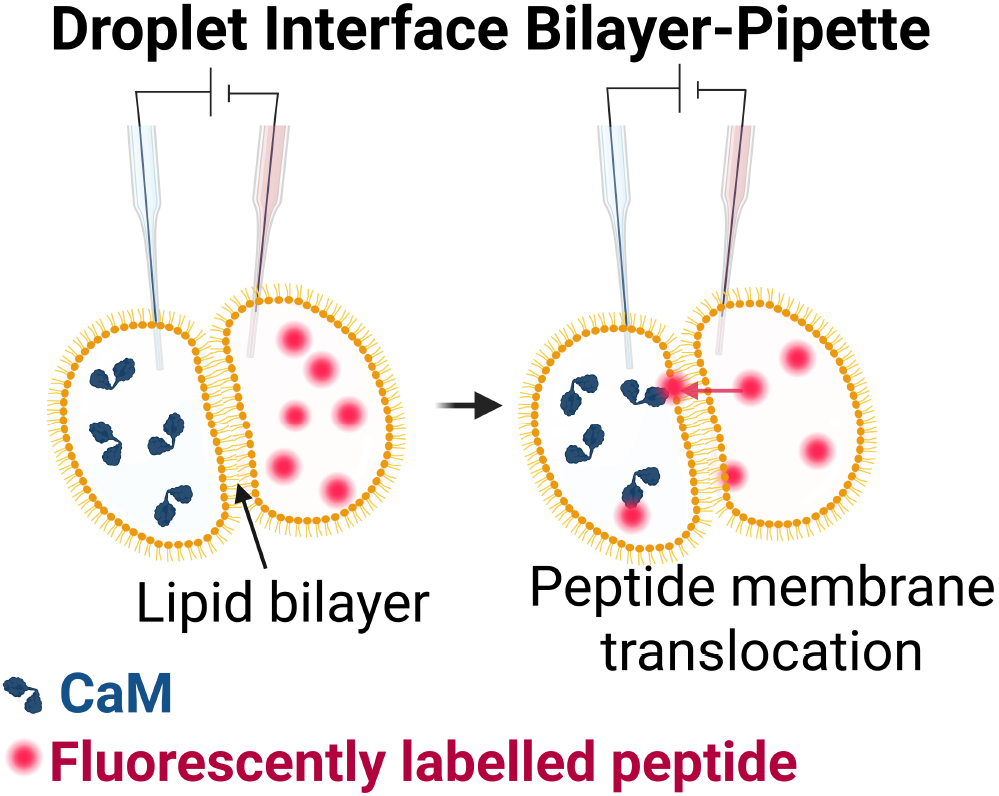

## Introduction

Whooping cough is a respiratory disease caused by *Bordetella pertussis*^1,2^, which remains particularly dangerous for unvaccinated individuals, especially newborns^3^. The adenylate cyclase (CyaA) toxin is one of the major virulence factors produced by *B. pertussis* and is involved in the early stages of respiratory tract colonization^4,5^. CyaA is a large, 1706-residue-long, multidomain protein composed of five domains (**Figure 1**): the N-terminal catalytic adenylate cyclase domain (ACD, residues 1–364)^6–8^, the translocation region (TR, residues 365–527)^9^, the hydrophobic region (HR, residues 528–710)^10,11^, the acylation region (AR, residues 711–1005)^9,12–19^, and the C-terminal Repeat-in-ToXin (RTX) receptor-binding domain (RD, residues 1006–1706)^15,20–30^. CyaA displays a unique intoxication mechanism in which ACD is directly translocated across the plasma membrane^31–33^ into the cytoplasm of target cells, where it produces supraphysiological levels of cAMP^7,28,31,34,35^, thereby subverting host defense^36–42^.

**Figure 1:**
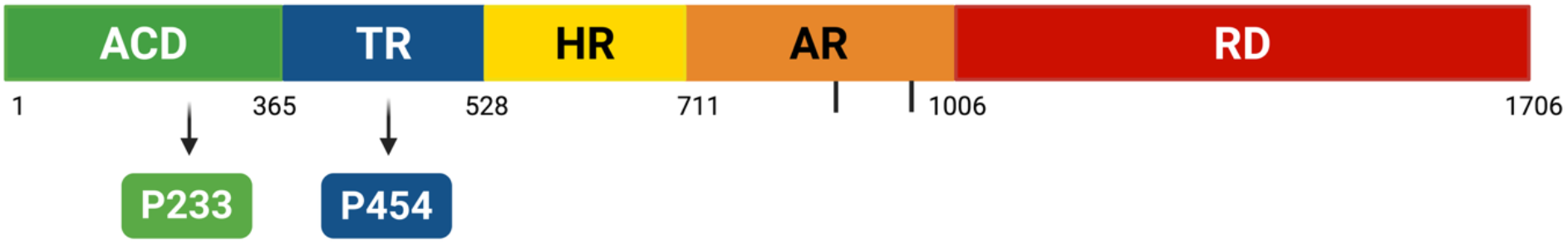
The CyaA toxin and the P233 and P454 peptides. Scheme of the CyaA toxin organization, showing its five domains: the N-terminal adenylate cyclase catalytic domain (ACD) in green, the translocation region (TR) in blue, the hydrophobic region (HR) in yellow, the acylation region (AR) in orange with the two acylations at K860 and K983 represented as black lines, and the RTX receptor binding domain (RD) in red. The segments P233 (from ACD) and P454 (from TR) are represented in green and blue, respectively. Made with Biorender.

Membrane translocation of ACD requires acylation of CyaA at two lysine residues within AR^16,43,44^ and depends on several host cell factors, including a negative membrane potential^16,45^, a calcium gradient across the plasma membrane^46^ and cytoplasmic calcium-loaded calmodulin (CaM)^16,47^. We previously showed that TR is essential to deliver ACD into the cytoplasm^9^ and that a peptide derived from this region, P454 (residues 454-484 of CyaA), exhibits membrane-active properties^48,49^. Membrane-active peptides are able to interact with membranes, eventually destabilizing them and/or translocating across them^50–54^. Typical membrane-active peptides are cell penetrating and antimicrobial peptides (CPPs and AMPs). Numerous AMPs kill bacteria by permeabilizing their membrane^55,56^, whereas CPPs can translocate across plasma membranes without permanently damaging it, preserving cell viability^57,58^. P454 interacts with membranes, translocates across membranes and binds CaM with high affinity^47–49^. In the context of CyaA intoxication, we previously showed that mutations within TR that reduce affinity for CaM correlate with a decrease in ACD translocation efficiency^47^. These observations led to the proposal that TR:CaM interactions generate an entropic pulling force that promotes ACD unfolding and membrane translocation^33,47^. Once in the cytoplasm, ACD binds CaM, which triggers its folding and enzymatic activation^7,8^. Formation of the ACD:CaM complex is primarily mediated by the H helix of ACD, a short, positively charged segment^6,8^. Interestingly, a peptide encompassing this region, P233 (residues 233-254 of CyaA), that binds CaM with high affinity, can also efficiently interact with membranes^8,47,49^.

Despite this qualitative and structural insights, the driving forces underlying ACD membrane translocation remain poorly understood. We previously showed that TR deletion from CyaA prevents ACD delivery into cells^9^ and that P454 exhibits an intrinsic ability to translocate across membranes and to bind CaM with high affinity^47^. These results led us to hypothesize that, once CyaA is inserted into the plasma membrane, TR translocates and binds CaM, thereby facilitating ACD translocation via a CaM-dependent entropic pulling mechanism^47^.

To further investigate these processes, we present here a new experimental approach called Droplet Interface Bilayer (DIB)-Pipette (**Figure 2**), derived from the DIB model membrane^59–64^, that enables to study membrane translocation events by epifluorescence microscopy while applying a controlled transmembrane potential to the lipid bilayer. Using this technique, we show that P454 can translocate across membranes both in the absence and in the presence of a transmembrane potential. Furthermore, the presence of positively charged residues within specific regions of ACD, such as the H helix (corresponding to the P233 sequence), led us to hypothesize that a physiological negative membrane potential may significantly contribute to their translocation. Consistent with this idea, we show that P233 does not translocate across membranes in the absence of a membrane potential but readily translocates when a negative potential is applied. Remarkably, we further show that the P233-P454 peptide, which corresponds to the sequence of the peptides P233 and P454 connected by a nine-residue-long glycine-serine linker, crosses membranes even in the absence of a membrane potential. Taken together, our results demonstrate that P233 and P454 exhibit distinct membrane-active properties, yet efficiently translocate across membranes, although in different conditions. These findings provide novel mechanistic insights into the driving forces governing ACD delivery into target cells.

**Figure 2:**
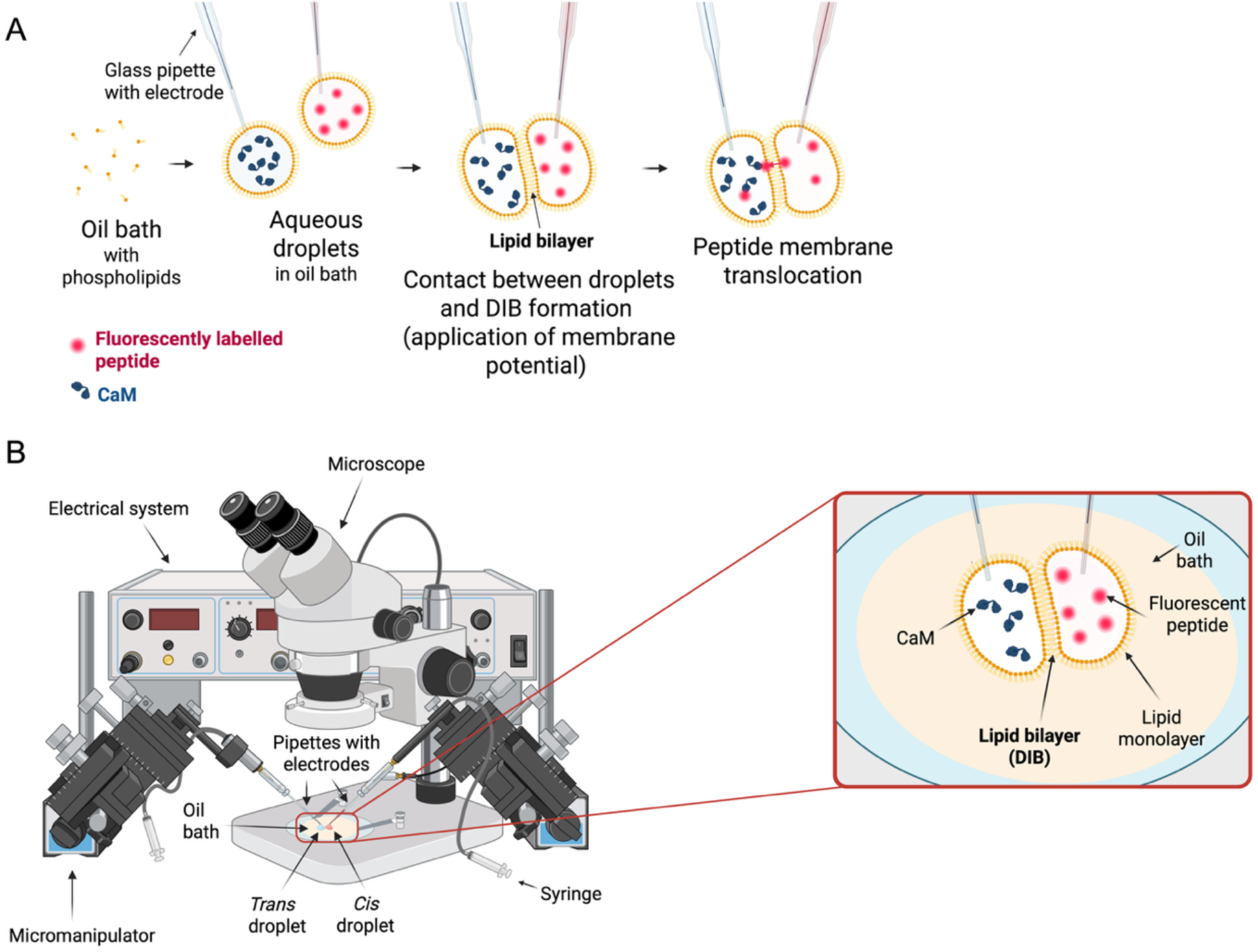
Schematic representation of the DIB-Pipette approach. **A)** Workflow of the DIB-Pipette approach: phospholipids are dissolved in an oil bath. Aqueous droplets, containing either fluorescently labelled peptide (*cis* droplet) or CaM (*trans* droplet), are injected into the oil bath using glass pipettes. Upon injection, a phospholipid monolayer assembles around each aqueous droplet. The droplets are brought into contact *via* pipette micromanipulation, leading to the formation of a lipid bilayer, termed Droplet Interface Bilayer, DIB. A membrane potential can be applied through electrodes inserted in the pipettes. Translocation of fluorescently labelled molecules across the bilayer may be observed either spontaneously or upon application of membrane potential, depending on the system. **B)** DIB-Pipette experimental set-up: the oil bath containing phospholipids is placed on a glass coverslip under a fluorescence microscope. Glass pipettes, filled with either fluorescently labelled peptide or CaM, are used to inject and manipulate *cis* and *trans* aqueous droplets within the oil bath. Droplet injection is achieved by gently applying air pressure through syringes connected to the pipettes. The pipettes are positioned and moved using micromanipulators, allowing precise control of droplets placement. Electrodes connected to an electrical system (composed of a digitizer and an amplifier) and inserted into the pipettes enable the application of a membrane potential on the system. Made with Biorender.

## Materials and methods

### Reagents

1,2-dioleoyl-sn-glycero-3-phosphocholine (DOPC, reference 850375C), 1-palmitoyl-2-oleoyl-sn-glycero-3-phosphocholine (POPC, reference 850457C), 1-palmitoyl-2-oleoyl-sn-glycero-3-[phospho-rac-(1-glycerol)] (POPG, reference 840457C), and cholesterol (ovine) (reference 700000P) were purchased from Avanti Polar Lipids. FITC-dextran 4.4 kDa was purchased from Sigma. Squalene was purchased from Sigma-Aldrich. Chloroform was purchased from Carlo Erba. All the experiments, unless stated otherwise, were performed in 20 mM HEPES, 150 mM NaCl, 2 mM CaCl_2_, pH 7.4 (buffer A).

### Peptides and proteins

All peptides were produced and purified by Genosphere Biotechnologies. Their purity and molecular masses were analyzed by HPLC and MALDI. Peptide P454 corresponds to residues 454 to 484 of CyaA and contains a single native tryptophan W458. Peptide P233 corresponds to the H helix in ACD, *i*.*e*., residues 233 to 254 of CyaA and contains the native tryptophan W242. Peptide P454-mut2 corresponds to residues 454 to 484 of CyaA (containing the native tryptophan W458) carrying four mutations: L463A-L475A-H477S-I479A. The P233-P454 peptide corresponds to the sequence of the peptides P233 and P454 connected by a nine-residue-long glycine-serine linker and contains the two native tryptophan residues W242 and W458 (see below for the primary sequences). All the peptides were also synthesized with a 5-carboxytetramethylrhodamine (TAMRA) linked to their N-terminus. The sequences of the peptides are the following:

- P454: [ac]-ASAHWGQRALQGAQAVAAAQRLVHAIALMTQ-[nh2]
- P233: [ac]-LDRERIDLLWKIARAGARSAVG-[nh2]
- P454-mut2: [ac]-ASAHWGQRAAQGAQAVAAAQRAVSAAALMTQ-[nh2]
- P233-P454: [ac]-LDRERIDLLWKIARAGARSAVGGGGSGG GGSASAHWGQRALQGAQAVAAAQRLVHAIALMTQ-[nh2]
- TAMRA-P454: [5Tamra]-ASAHWGQRALQGAQAVAAAQRLVHAIALMTQ-[nh2]
- TAMRA-P233: [5Tamra]- LDRERIDLLWKIARAGARSAVG-[nh2]
- TAMRA-P454-mut2: [5Tamra]- ASAHWGQRAAQGAQAVAAAQRAVSAAALMTQ-[nh2]
- TAMRA-P233-P454: [5Tamra]- LDRERIDLLWKIARAGARSAVGGGGSGG GGSASAHWGQRALQGAQAVAAAQRLVHA IALMTQ-[nh2]

Human calmodulin (CaM) was produced in *E. coli* and purified as described in ^47^. Briefly, CaM was precipitated with ammonium sulphate followed by glacial acetic acid precipitation. CaM was subsequently purified by a hydrophobic interaction chromatography on Phenyl Sepharose (EDTA-CaM), an ion-exchange chromatography on Q-Sepharose fast flow, a second hydrophobic interaction chromatography on Phenyl Sepharose (calcium-CaM) and a final size exclusion chromatography on a Sephacryl S100.

### Lipid vesicles preparation

Small unilamellar vesicles (SUVs) were prepared at a final lipid concentration of 40 mM. The following membrane compositions were prepared: POPC 100%, POPC:POPG 8:2, and DOPC:POPG 8:2 (in HEPES 20 mM, NaCl 150 mM, pH 7.4), as previously described^48,49^. Briefly, MLVs (multilamellar vesicles) were prepared by reverse phase evaporation, passed through polycarbonate filters with 1.2 μm diameter (Merck Millipore) and subsequently sonicated using a tip sonicator to obtain SUVs. SUVs hydrodynamic diameters, dispersity, and charge were checked by dynamic light scattering (DLS) and electrophoretic mobility using a NanoZS instrument (Malvern Instruments).

### Membrane partitioning followed by B2LiVe

B2LiVe NMR experiments were performed as described in ^65^. Experiments were performed on a Bruker Avance NEO 800 MHz (Bruker Biospin, Billerica, USA) with an 18.8 Tesla magnetic field, equipped with a cryogenically cooled triple resonance TCI probe. The samples used for NMR and tryptophan fluorescence experiments were derived from a common stock solution to ensure comparability between the two methods. P233 concentration was 5 μM for all samples while lipid concentration was increased from 0 up to 10 mM. Titrations were performed at 25°C in the presence of SUVs composed of POPC:POPG in a molar ratio of 8:2, DOPC:POPG in a molar ratio of 8:2 and POPC 100% in buffer A, supplemented with 5% of D_2_O (Eurisotop, Saclay, France).

### Membrane partitioning followed by tryptophan fluorescence

The samples used for NMR and tryptophan fluorescence experiments were derived from a common stock solution to ensure comparability between the two methods. Fluorescence titrations were performed with a FP-8200 Jasco spectrofluorometer, equipped with a Peltier-thermostat ETC-272T (25°C). A 5 nm bandwidth was used for both excitation and emission. Fluorescence experiments were performed in a 105.251.QS cuvette (Hellma). Fluorescence emission spectra were recorded from 300 to 400 nm at a scan rate of 100 nm/min, using an excitation wavelength of 280 nm. Fluorescence emission spectra of the peptides were corrected for SUVs light scattering. The wavelength of maximal fluorescence emission (λmax) and fluorescence intensity ratio at 340 nm over 380 nm were used to report peptide-membrane interaction and to measure the partition coefficient *K*_*x*_. P233 concentration was 5 μM for all samples, while lipid concentration was increased from 0 up to 10 mM. Titrations were performed at 25°C in the presence of SUVs composed of POPC:POPG in a molar ratio of 8:2, DOPC:POPG in a molar ratio of 8:2 and POPC 100% in buffer A, supplemented with 5% of D_2_O (Eurisotop, Saclay, France).

### Determination of dissociation constant *K*_*d*_, partition coefficient *K*_*x*_ and free energy of partitioning *ΔG*

In this work we refer to “solution to membrane partitioning” or simply “membrane partitioning” as the process by which a molecule of interest (*e*.*g*. a peptide) distributes, between the aqueous phase and the lipid bilayer environment. At equilibrium, the distribution of an amphitropic peptide is quantified by the partition coefficient (*K*_*x*_) on a mole-fraction scale. The solution to membrane partitioning coefficient, *K*_*x*_, the dissociation constant *K*_*d*_, and the free energy of partitioning *ΔG*_*Kx*_ were determined by analysis of the fluorescence intensity ratios at 340/380 nm and of 1D-NMR data.

The partition coefficient *K*_*x*_ is defined as the ratio of the protein concentrations in the lipid (P_L_) environment and in water (P_W_) phases^49^. It is expressed as follows:

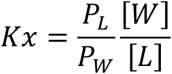

where [*W*] represents the water concentration, and [*L*] the lipid concentration.

The fraction of peptide partitioned into the membrane phase, *f*_*PL*_, is given by equation (1):

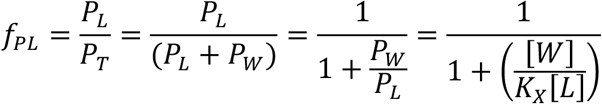

The partition coefficient is related to the apparent dissociation constant as follows: *K*_*x*_ × *K*_*d*_ = [*W*] with *K*_*d*_ × *P*_*L*_ = *P*_*W*_ × [*L*]. The equation *f*_*PL*_ is fitted to the experimental data using KaleidaGraph (Synergy Software), providing *K*_*x*_ and *K*_*d*_. In NMR experiments, an increase of signal intensity at low lipid concentrations corresponding to transient interactions of P233 was observed. The data was fit to a sum of equations (1) and only the parameters corresponding to the signal loss (expected for membrane partitioning) were considered to determine the membrane partitioning constants *K*_*x*_ and *K*_*d*_.

Once the partition coefficient is calculated, the free energy *(ΔG*) of transfer of the peptide from water to membrane was calculated as follows *ΔG = RT × ln (K*_*x*_), where *ΔG* is the free energy (kcal·mol^-1^), *R* is the gas constant (R = 1.98 10^-3^ kcal·mol^-1^·K^-1^), *T* is the temperature in Kelvin, and *K*_*x*_ is the partition coefficient.

### Far-UV Synchrotron radiation circular dichroism

Far-UV synchrotron radiation circular dichroism (SRCD) experiments were performed at the DISCO beamline of the Synchrotron SOLEIL (Gif-sur-Yvette). SRCD spectra were recorded at 25 °C with an integration time of 1.2 s and a bandwidth of 1 nm with a resolution of 1 nm. Each far-UV spectrum represents the average of four individual scans. QS cells (Hellma) with a pathlength of 100 μm were used to record the SRCD signal in the far-UV from 190 to 250 nm. Peptides P233 and P454 were scanned at a concentration of 100 μM, peptide P454-mut2 at 130 μM and P233-P454 at 80 μM in buffer A in the absence and presence of 40% trifluoroethanol (TFE). Peptides were also scanned in presence of SUVs (POPC, POPC:POPG 8:2, or DOPC:POPG 8:2) at final concentration of 6 mM lipids. Buffer A, in the absence or presence of SUVs and 40% TFE, was used as blank and subtracted from the far-UV CD spectra of the peptides. Secondary structure content was estimated using BestSel^66^.

### Synchrotron radiation oriented-circular dichroism (SR O-CD)

Synchrotron radiation oriented-circular dichroism (SR O-CD) experiments were performed at the DISCO beamline of the Synchrotron SOLEIL (Gif-sur-Yvette).

Supported lipid bilayer preparation: cells were washed successively with Hellmanex^®^ 2%, ethanol and water, and then dried under nitrogen flow. They were successively sonicated in isopropanol and MilliQ water baths three times. After drying, cells were ozone-cleaned using “UV Ozone Cleaner – ProCleaner Plus” (Bioforce Nanosciences) for 20 minutes, rinsed with isopropanol, Milli-Q water, and isopropanol again. Cells were then dried under nitrogen flow. 70 µL of freshly sonicated SUVs at a lipid concentration of 150 µM in buffer A were brought into contact with the silica surface for 15 minutes. Non-fused SUVs were rinsed with 10 mL of Milli-Q water and buffer A.

Peptide deposition: the solution containing 25 µg of peptide solubilized in 80 μL MilliQ water was deposited on the newly formed SLB. The water was removed under vacuum for 1 hour. Subsequently, cells were mounted into the measurement chamber at 98% humidity (maintained by a K_2_SO_4_ saturated solution) at 25°C and incubated in the humidity chamber for at least 1 hour. Temperature was controlled using a JULABO recirculating thermostatic bath. After measurement, any dissociated proteins/peptides from SLB could be rinsed away. Washing steps consist of meticulous washing with 10 mL of buffer. Excess volume of buffer was then removed under vacuum, and the cell was rehydrated as described before.

O-CD measurements: circular dichroism (CD) spectra were recorded at a scan rate of 30 nm.min^-1^ with measurements taken at 8 distinct angles (rotated by 45° increments). Spectra were acquired in duplicated at each angle. Linear dichroism spectra were acquired at the end of the measurement at a single angle to assess the viability of the SLB. After acquisition, the 16 CD measurements were averaged, normalized, and baseline-corrected to zero between 250 and 255 nm.

### ANTS-DPX permeabilization assays

ANTS (fluorophore probe) and DPX (quencher) were encapsulated into large unilamellar vesicles (LUVs) to monitor membrane permeabilization induced by the peptides. LUVs were prepared at a lipid concentration of 20 mM (at the following molar ratio: DOPC:POPG 8:2) in HEPES 20 mM, NaCl 150 mM pH 7.4 containing 40 mM ANTS and 120 mM DPX. LUVs were prepared from MLVs (obtained as previously described for the SUVs preparation). The MLVs suspension was passed through 1.2, 0.8, 0.4 and 0.2 μm polycarbonate filters. The unencapsulated probes were removed by gel filtration using a Sephadex G-25 column (5 mL) (GE Healthcare Life Sciences, Pittsburgh, PA, USA). For permeabilization assays, LUVs were incubated in buffer A at 0.4 mM lipids at 25°C in a 96-well plate (ThermoFisher Scientific) in the presence of various peptide concentrations ranging from 0.1 to 10 μM. The excitation wavelength was set to 360 nm and the fluorescence of ANTS was continuously monitored at 520 nm for 1 hour with a FP-5200 Jasco spectrofluorometer. The percentage of fluorescence intensity was calculated using the maximum intensity measured after complete disruption of the vesicles achieved by the addition of 0.12% (v/v) Triton X100 (2 mM).

### Droplet interface bilayer (DIB) pipette method

An oil bath was prepared by dissolving 15% or 20% (v/v) chloroform and 0.08% or 0.1% (w/v) lipids (DOPC:POPG at an 8:2 molar ratio) in squalene. Two aqueous buffers were prepared. The *cis* buffer is composed of 20 mM HEPES, 150 mM NaCl, 8 mM MgCl_2_, 2 mM CaCl_2_, pH 7.4 and the *trans* buffer is composed of 20 mM HEPES, 150 mM NaCl, 9.8 mM MgCl_2_, 0.2 mM CaCl_2_, pH 7.4. Consequently, two aqueous solutions were prepared: the *cis* solution containing 10 μM of fluorescently labelled peptides/FITC-dextran in *cis* buffer and the *trans* solution containing 10 μM of CaM in *trans* buffer. We previously showed that CaM is required to extract the peptide once translocated at the membrane surface of the *trans* droplet to ensure the accumulation of peptide in solution and facilitate its quantification by fluorescence in the lumen of the *trans* droplet. Indeed, in the absence of CaM, the affinities of P233 and P454 for membranes prevents membrane-to-solution partitioning of the peptides. The formation of a fluorescent ring at the surface of the *trans* droplet lipid monolayer does not allow fluorescence quantification^47^.

Glass pipettes were pulled from glass capillaries (Harvard apparatus) using a pipette puller (Nashirige PC-100). Silver electrodes cut from a silver wire (Phymep) were pretreated by overnight incubation in bleach followed by extensive rinsing with Milli-Q water prior to use. *Cis* and *trans* pipettes were respectively filled with the *cis* and *trans* solutions. Pipettes were then mounted on micromanipulators (Sutter instruments), connected to syringes and were consequently immersed in dimethyldichlorosilane (DMDCS, 5% in ethanol) for three minutes and rinsed in ethanol for one minute to ensure proper droplet adhesion. Oil bath (100 µL) was deposited onto a glass coverslip covered with a thin layer of polydimethylsiloxane, PDMS (Sylgard). Pipettes were immersed in the oil bath by micromanipulation. Droplets were injected from the pipettes into the oil bath by gently applying air pressure from two syringes connected to the glass pipettes (**Figure 2**) and maintained separate for one to three minutes before being brought into contact *via* micromanipulation. Once in close proximity, a lipid bilayer (the DIB) is formed at the interface between the two droplets. Upon DIB formation, a membrane potential of either -80 or -150 mV was applied when required. The membrane potential was imposed using an amplifier (Axopatch 200B) and a digitizer (Digidata 1440A), *via* the silver electrodes inserted into the glass pipettes. The electric signal was monitored using the pClamp10 software. Potential translocation of the fluorescently labelled peptides and FITC-dextran across the DIB was monitored by fluorescence microscopy for 15 minutes using an inverted microscope (Olympus IX71) and a camera (uEye, IDS UI-3060CP-M-GL REV.2). Fluorescent probes were excited using a mercury lamp and excitation filters (515-555 nm for TAMRA-labelled peptides, 420-560 nm for the FITC-dextran).

### Quantification of peptides/FITC-dextran translocation

The translocation process was quantified by analyzing fluorescence images of droplet pairs using ImageJ-4. Since fluorescence intensity is not strictly proportional to peptide concentrations - due to photobleaching, small volume variations, peptides leakage from droplets and peptide adsorption to glass pipettes - relative change in fluorescence between *cis* and *trans* droplets were used to assess translocation. For each image, fluorescence intensities were measured by sampling three circular regions of interest (ROIs) in the *cis* droplet, *trans* droplet, and background. Background fluorescence was subtracted from both *cis* and *trans* droplet fluorescence. The amount of peptide reaching the *trans* droplet was estimated by calculating the ratio of *trans* to *cis* fluorescence, after background subtraction. The effective increase in fluorescence at a given time was determined by subtracting the initial fluorescence ratio at time 0 as follows: ΔFluorescence ratio = Fluorescence ratio_t=x_ – Fluorescence ratio_t=0_. Replicates were averaged. The error bars correspond to the errors of the averages calculated based on the errors of the subtractions of the fluorescence ratios at time t=x and t=0. Translocation is considered to occur strongly if ΔFluorescence ratio ≥ 5%, to occur weakly if 5% > ΔFluorescence ratio > 2% and is considered absent if ΔFluorescence ratio ≥ 2%. These thresholds have been set taking into consideration that the average ΔFluorescence ratio of FITC-dextran at t=15 minutes is ∼ 0% and the average standard deviation in ratio determination (σ) registered during translocation events is ∼ 1%. Therefore, the threshold to consider that translocation does not occur has been set at 2σ (ΔFluorescence ratio = 2%) and the threshold to indicate significant, unambiguous translocation events has been set at 5σ (ΔFluorescence ratio = 5%). Different experimental conditions were compared using an adaptation for 3 possible ordered results of the Fisher’s exact test. Results were considered statistically significant for *p value* ≤ 0.05.

### Schematic figures

The schematic figures were created using BioRender.

## Results

### P454 is a membrane-active peptide able to cross membranes

We have previously reported that the P454 peptide exhibits membrane-active properties^47–49^. In line with these results, here, we show by far-UV SRCD that P454 is mainly unstructured in solution, and folds into helical structures upon membrane interaction (**Figure 3A**). Moreover, we show that P454 permeabilizes lipid vesicles (**Figure 3C**). We previously reported that the fluorescently labelled P454 peptide, TAMRA-P454, was able to cross a lipid bilayer in the absence of membrane potential, in the presence of calcium-loaded calmodulin (CaM) in the *trans* compartment. CaM binds and extracts P454 from the *trans* side of the lipid bilayer, facilitating quantification of the peptide in the *trans* compartment of the membrane. In the absence of CaM, P454 remains predominantly localized at the lipid–buffer interface of the *trans* compartment^47^. Here, we investigate the membrane translocation of TAMRA-P454 by means of our newly developed DIB approach, called DIB-Pipette. Unlike the approach we previously used^47^, which was based on the formation of random droplet pairs by emulsion mixing, the DIB-Pipette approach enables a controlled droplet formation and a precise bilayer assembly by means of glass pipettes. The presence of electrodes, inserted in the glass pipettes, allows direct application of a membrane electric potential to the lipid bilayer, which is essential for investigating voltage-dependent translocation mechanisms. Moreover, the presence of a single droplet pair strongly improves the signal-to-noise ratio, thanks to the reduced background fluorescence (**Figure 2**). The results indicate that TAMRA-P454 can translocate across membranes composed of DOPC:POPG (at a 8:2 molar ratio) both in the absence and in the presence of a negative membrane potential (-150 mV) (**Figure 3D**). These results confirm the membrane-active properties of P454 and its ability to cross membranes independently of any membrane potential.

**Figure 3:**
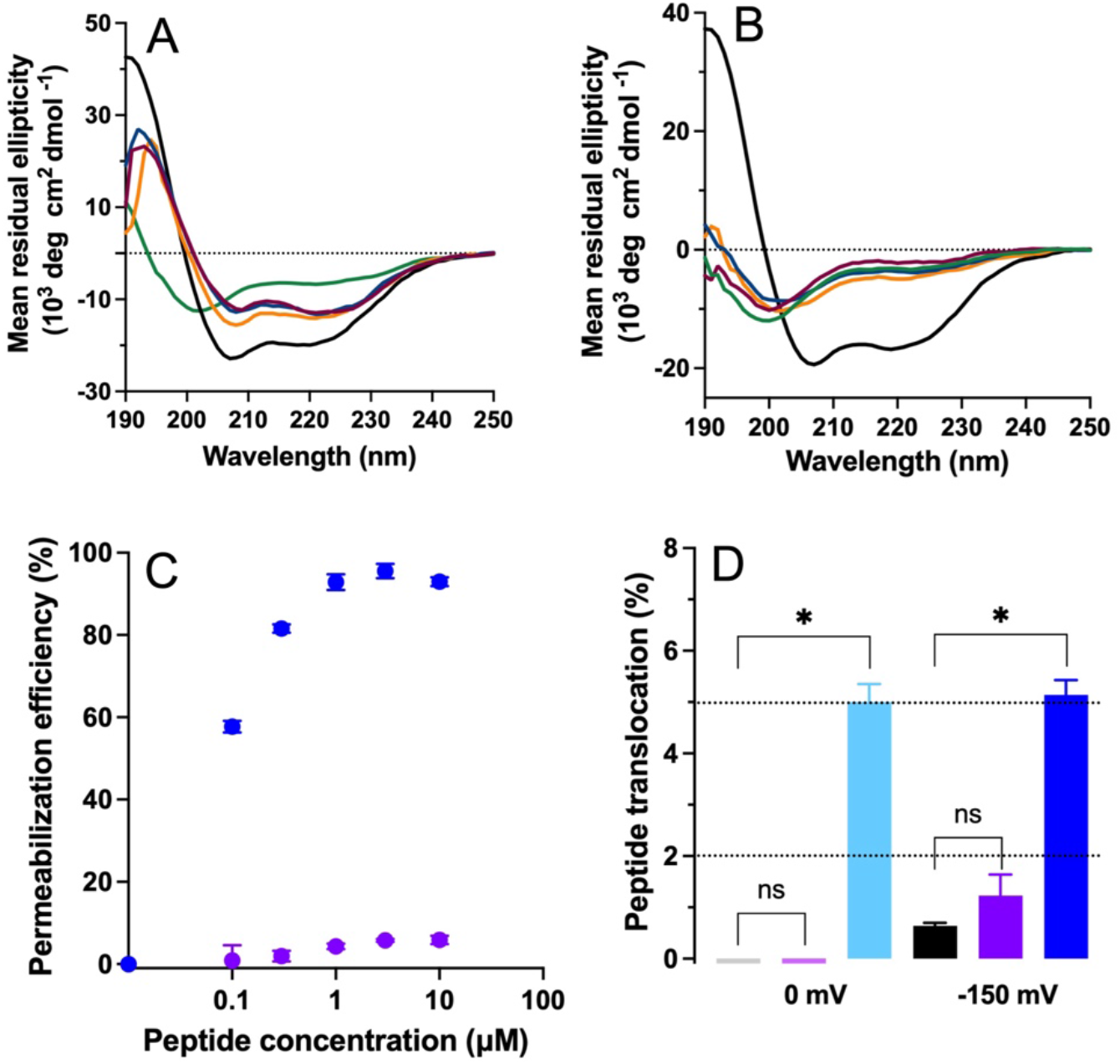
Measurement of the membrane-active properties of P454 and P454-mut2. Far-UV synchrotron radiation circular dichroism (SRCD) spectra of **A)** 100 µM P454 and **B)** 130 µM P454-mut2 recorded in the absence (green line) and in the presence of 6 mM SUVs composed of POPC (burgundy line), POPC:POPG at an 8:2 molar ratio (blue line), or DOPC:POPG at an 8:2 molar ratio (orange line), as well as in the presence of 40% TFE (black line) in buffer A at 25 °C. **C)** Membrane permeabilization measured as percentage of efflux of the fluorophore probe ANTS and its quencher DPX from DOPC:POPG (8:2 molar ratio) large unilamellar vesicles (LUVs; 0.4 mM lipids) induced by P454 (in blue) and P454-mut2 (in violet) after 30 minutes of incubation at 25°C at indicated peptide concentrations (0.1–10 µM). **D)** Membrane translocation of TAMRA-P454 and TAMRA-P454-mut2 in the absence (light blue and light violet, respectively) and presence (dark blue and dark violet, respectively) of a −150 mV membrane potential measured by the DIB-Pipette approach. Membrane translocation is expressed as the increase (percentage) in the ratio between fluorescence in the *trans* droplet and fluorescence in the *cis* droplet after 15 minutes of incubation. TAMRA-P454 and TAMRA-P454-mut2 translocation is compared with that of a negative control molecule (FITC-dextran) in the absence (grey) and presence (black) of a −150 mV membrane potential. Black dotted lines at y = 2% and y = 5% indicate the thresholds for translocation as described in the **Materials and Methods** section.

We previously showed that the substitution of four residues (L463A, L475A, H477S, I479A) within the P454 sequence in the context of the full-length CyaA toxin completely abolishes the toxin’s ability to intoxicate cells. We also reported that a peptide carrying the same point mutations had a reduced affinity for CaM and for membranes^47^. Here, we show that this mutant peptide, called P454-mut2, while being unstructured in solution like the wild type P454, does not form helical structures in the presence of membranes (**Figure 3B**) and completely loses its ability to permeabilize lipid vesicles (**Figure 3C**). Compared to the wild type P454 peptide, the DIB-Pipette experiments (**Figure 3D**) show that TAMRA-P454-mut2 is unable to translocate across membranes both in the absence and in the presence of a negative membrane potential of -150 mV. Thus, the L463A, L475A, H477S, I479A mutations abrogate the membrane-active properties of the P454 region without affecting the mean net charge of the peptide.

### The P233 peptide requires an electric membrane potential to cross membranes

The P233 peptide, which encompasses the main CaM binding site of ACD^6,8^, is also able to interact with membranes^49^. In this work, we further investigated P233 membrane interactions by characterizing its affinity for membranes composed of either POPC, POPC:POPG 8:2 molar ratio, or DOPC:POPG 8:2 molar ratio. Membrane affinity was assessed using two complementary approaches: B2LiVe, a NMR-based method^65^, and tryptophan fluorescence. Both techniques indicate that P233 interacts with SUVs (**Figure 4A and Figure 4B, Figure S1 - Supplementary Information**). The partition coefficient (*K*_*x*_) and free energy of solution to membrane partitioning (*ΔG*_*Kx*_) of P233 were estimated from the tryptophan intrinsic fluorescence titration curves (**Figure 4A**) and the NMR spectra (**Figure 4B**) as described in the **Materials and methods** section. The *K*_*x*_ values reported (**Table S1 - Supplementary Information**) indicate that P233 has a rather low affinity for SUVs containing only zwitterionic lipids. However, in the presence of negatively charged lipids (20% PG), the affinity of P233 for membranes increases, even more when POPC is substituted with DOPC (**Table S1 - Supplementary Information**), indicating that negative charges and membrane fluidity increase P233 affinity for membranes. We also measured the secondary structure content of the P233 peptide in the absence and presence of membranes of various lipid compositions by far-UV SRCD (**Figure 4C**): our results indicate that the peptide is mainly unstructured in solution but forms helical structures in the presence of SUVs containing 20% POPG. Moreover, as reported by SR O-CD (Synchrotron Radiation Oriented-Circular Dichroism) experiments, the helical structures formed by P233 in the presence of negatively charged membranes are tilted with respect to the plane of the bilayer (**Figure 4D**). Finally, we showed that P233 is able to permeabilize lipid vesicles in our experimental conditions (**Figure 4E**), although less efficiently than P454 (**Figure 3C**). Taken together, our results indicate that P233 exhibits membrane-active properties and prompt the question of whether P233 could also translocate across membranes. We previously reported that fluorescently-labelled P233, TAMRA-P233, was not able to translocate across a lipid bilayer in the absence of membrane potential^47^. We confirmed this result using the DIB-Pipette approach (**Figure 4F**). However, the presence of positively charged residues within the P233 sequence led us to hypothesize that a negative membrane potential may trigger P233 membrane translocation. Therefore, using the DIB-Pipette approach, we applied a negative membrane potential (-150 mV) to the system and evaluated TAMRA-P233 translocation. Our results indicate that the presence of a negative membrane potential triggers TAMRA-P233 translocation (**Figure 4F**). We repeated these experiments in the presence of a more physiological membrane potential^66^ (-80 mV) and found that the TAMRA-P233 peptide can efficiently translocate across membranes in these conditions (**Figure 4F**).

**Figure 4:**
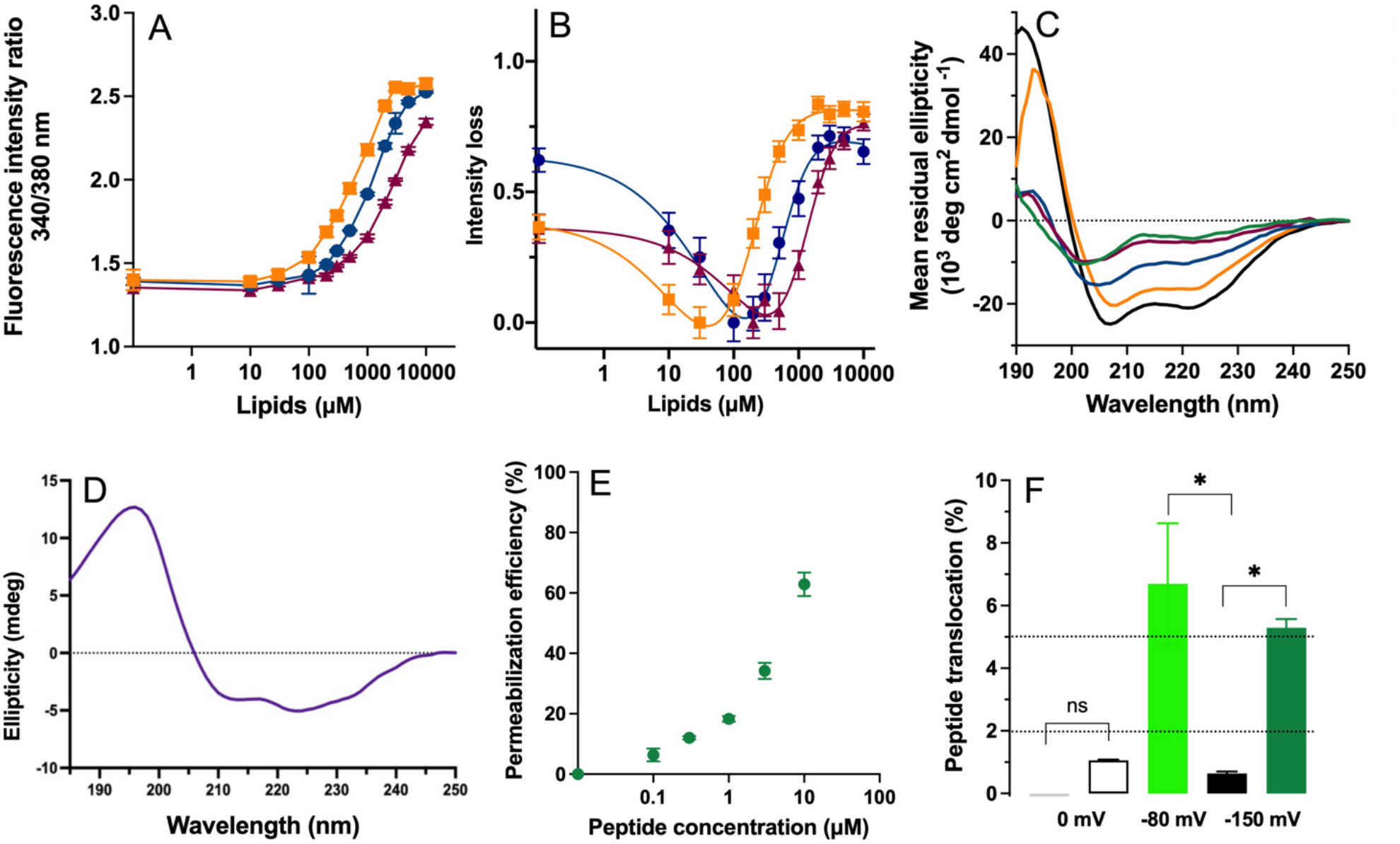
Measurement of the membrane-active properties of P233. Membrane partitioning followed by **A)** tryptophan fluorescence or **B)** by B2LiVe of P233 (5 µM) with SUVs composed of POPC (burgundy triangles), POPC:POPG at an 8:2 molar ratio (blue circles), or DOPC:POPG at an 8:2 molar ratio (orange squares) at 25°C at increasing lipid concentrations (10-10,000 µM). Solid lines correspond to fits obtained as described in **Materials and Methods. C)** Far UV synchrotron radiation circular dichroism (SRCD) spectra of 100 µM P233 recorded in the absence (green line) and in the presence of 6 mM SUVs composed of POPC (burgundy line), POPC:POPG at an 8:2 molar ratio (blue line), or DOPC:POPG at an 8:2 molar ratio (orange line), as well as in the presence of 40% TFE (black line) in buffer A at 25 °C. **D)** Synchrotron radiation oriented-circular dichroism (SR O-CD) spectra of P233 recorded on POPC:POPG 8:2 molar ratio supported lipid bilayers (SLBs). **E)** Membrane permeabilization measured as percentage of efflux of the fluorophore probe ANTS and its quencher DPX from DOPC:POPG (8:2 molar ratio) large unilamellar vesicles (LUVs; 0.4 mM lipids) induced by P233 after 30 minutes of incubation at 25 °C at indicated peptide concentrations (0.1–10 µM). **F)** Membrane translocation of TAMRA-P233 determined by the DIB-Pipette approach in the absence (white) and presence of a membrane potential of either - 80 mV (light green) or -150 mV (dark green). Membrane translocation is expressed as the increase (percentage) in the ratio between fluorescence in the *trans* droplet and fluorescence in the *cis* droplet after 15 minutes of incubation. TAMRA-P233 translocation is compared with that of a negative control molecule (FITC-dextran) in the absence (grey) and presence (black) of a −150 mV membrane potential. Black dotted lines at y = 2% and y = 5% indicate the thresholds for translocation as described in the **Materials and Methods** section.

### The P233-P454 peptide translocates across membranes in the absence of a membrane potential

Recent works indicate that couples of different peptides are able to synergistically potentialize their membrane-active properties^68,69^. Thus, we analyzed the potential synergistic activity of P233 and P454 peptides to favor membrane translocation of P233 in the absence of a membrane potential. The DIB-Pipette assays indicate that co-incubation of TAMRA-P233 and non-labelled P454 is not sufficient to promote TAMRA-P233 translocation across membranes in the absence of a membrane potential (**Figure 5**). However, the membrane-active properties of P454^32,47,48^ lead to the hypothesis that it could drive the translocation of other CyaA segments. To test this hypothesis, we designed a peptide, P233-P454, that contains both P233 and P454 sequences connected by a nine-residue-long glycine and serine linker. As evidenced by far-UV SRCD, P233-P454 is mainly unstructured in solution and forms helical structures upon membrane interaction (**Figure 6A**), similarly to the isolated P233 and P454 peptides (**Figure S2 - Supplementary Information**). Moreover, permeabilization assays indicated that P233-P454 efficiently permeabilizes membranes (**Figure 6B**). Noteworthy, the DIB-Pipette experiments show that TAMRA-P233-P454 is able to translocate across membranes in the absence of a membrane potential, as reported by the time-dependent fluorescence increase reported in the *trans* droplet (**Figure 6C**). This result shows that, in the absence of a membrane potential, P454 drives P233 membrane translocation when the two peptides are covalently linked. In contrast, isolated P233 crosses membranes only in the presence of a membrane potential (**Figure 4F**). Taken together, our results indicate that P454 possesses intrinsic membrane-translocation capability, whereas P233 requires either a membrane potential or the energy provided by the P454:CaM translocation complex to cross membranes.

**Figure 5:**
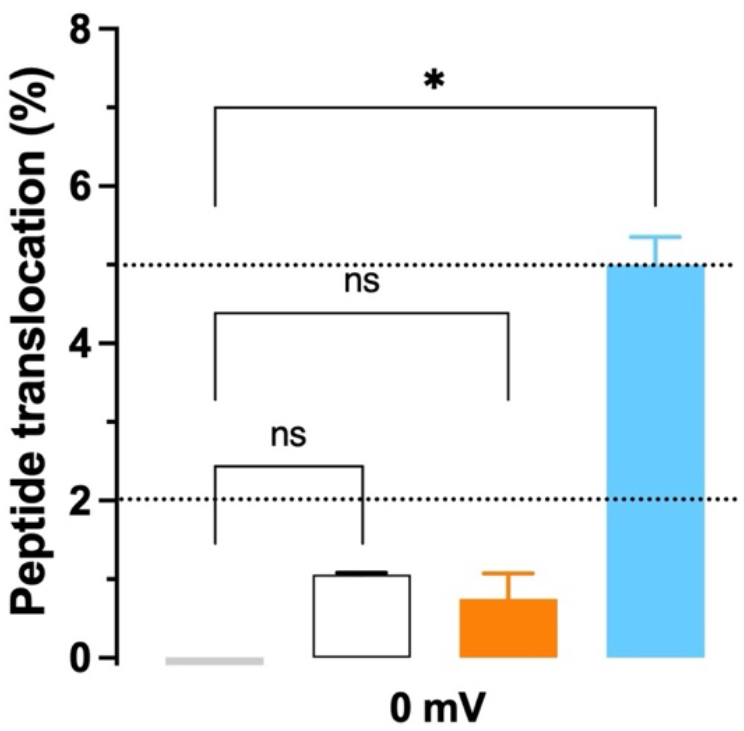
Measurement of TAMRA-P233 membrane translocation in the presence of non-labelled P454. Membrane translocation of TAMRA-P233 determined by the DIB-Pipette approach in the absence (in white) or presence (in orange) of non-labelled P454. Membrane translocation is expressed as the increase (percentage) in the ratio between fluorescence in the *trans* droplet and fluorescence in the *cis* droplet after 15 minutes of incubation. TAMRA-P233 translocation in the absence and presence of P454 is compared with that of a negative control molecule (FITC-dextran) (in grey) and that of TAMRA-P454 (in light blue). Black dotted lines at y = 2% and y = 5% indicate the thresholds for translocation as described in the **Materials and Methods** section.

**Figure 6:**
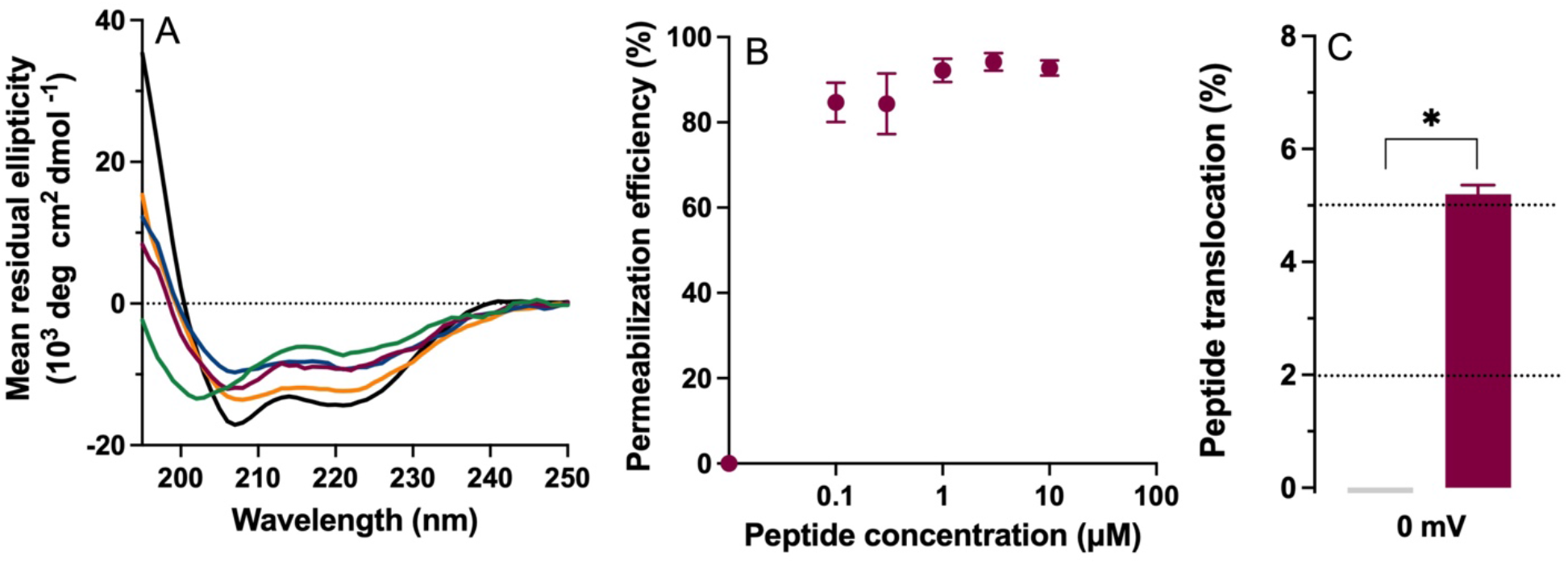
Measurement of the membrane-active properties of P233-P454. **A)** Far UV synchrotron radiation circular dichroism (SRCD) spectra of 80 µM P233-P454 recorded in the absence (green line) and in the presence of 6 mM SUVs composed of POPC (burgundy line), POPC:POPG at an 8:2 molar ratio (blue line), or DOPC:POPG at an 8:2 molar ratio (orange line), as well as in the presence of 40% TFE (black line) in buffer A at 25 °C. **B)** Membrane permeabilization percentage measured as efflux of the fluorophore probe ANTS and its quencher DPX from DOPC:POPG (8:2 molar ratio) large unilamellar vesicles (LUVs; 0.4 mM lipids) induced by P233-P454 (in burgundy) after 30 minutes of incubation at 25 °C at indicated peptide concentrations (0.1–10 µM). **C)** Membrane translocation of TAMRA-P233-P454 (in burgundy) determined by the DIB-Pipette method in the absence of a membrane potential. Membrane translocation is expressed as the increase (percentage) in the ratio between fluorescence in the *trans* droplet and fluorescence in the *cis* droplet after 15 minutes of incubation. TAMRA-P233-P454 translocation is compared with that of a negative control molecule (FITC-dextran) (in grey) in the absence of a membrane potential. Black dotted lines at y = 2% and y = 5% indicate the thresholds for translocation as described in the **Materials and Methods** section.

## Discussion

### P233 and P454 have different membrane-active properties

In this work, we investigated the membrane translocation mechanism of the *Bordetella pertussis* adenylate cyclase (CyaA) toxin, focusing on the functional contributions of two short peptide segments, P233 from the adenylate cyclase catalytic domain (ACD), and P454 from the translocation region (TR). In the presence of calcium, both peptides interact with high affinity with the endogenous eukaryotic protein calmodulin (CaM), to promote efficient translocation of the CyaA ACD into the host cell cytoplasm^47^. Using a recently developed Droplet Interface Bilayer (DIB) approach called DIB-Pipette, that enables direct visualization of fluorescently labelled molecules transport under controlled transmembrane potentials, we showed that these two peptides exhibit fundamentally distinct translocation behaviors (**Figure 7**). While P454 spontaneously translocates across lipid bilayers independently of membrane potential, P233 requires a negative membrane potential to do so. Notably, covalent coupling of P233 to P454 enables efficient translocation of P233 even in the absence of a membrane potential. Although both P454 and P233 interact with lipid membranes, fold into helical conformations upon membrane interaction, and permeabilize lipid bilayers, their membrane-active properties, as well as their functional roles in CyaA translocation differ sharply. P454 behaves as an autonomous membrane-translocating segment, capable of overcoming the energetic cost of membrane translocation without the need of an electric field. In contrast, P233, despite efficient membrane interaction, remains unable to translocate across membranes unless a negative transmembrane potential is applied. This observation indicates that membrane insertion and membrane translocation are not equivalent processes. Rather, P233 appears to populate interfacial or partially membrane-inserted states that are separated from the translocated state by an energy barrier that cannot be overcome by thermal fluctuations alone and instead requires either a transmembrane potential or covalent coupling to P454.

**Figure 7:**
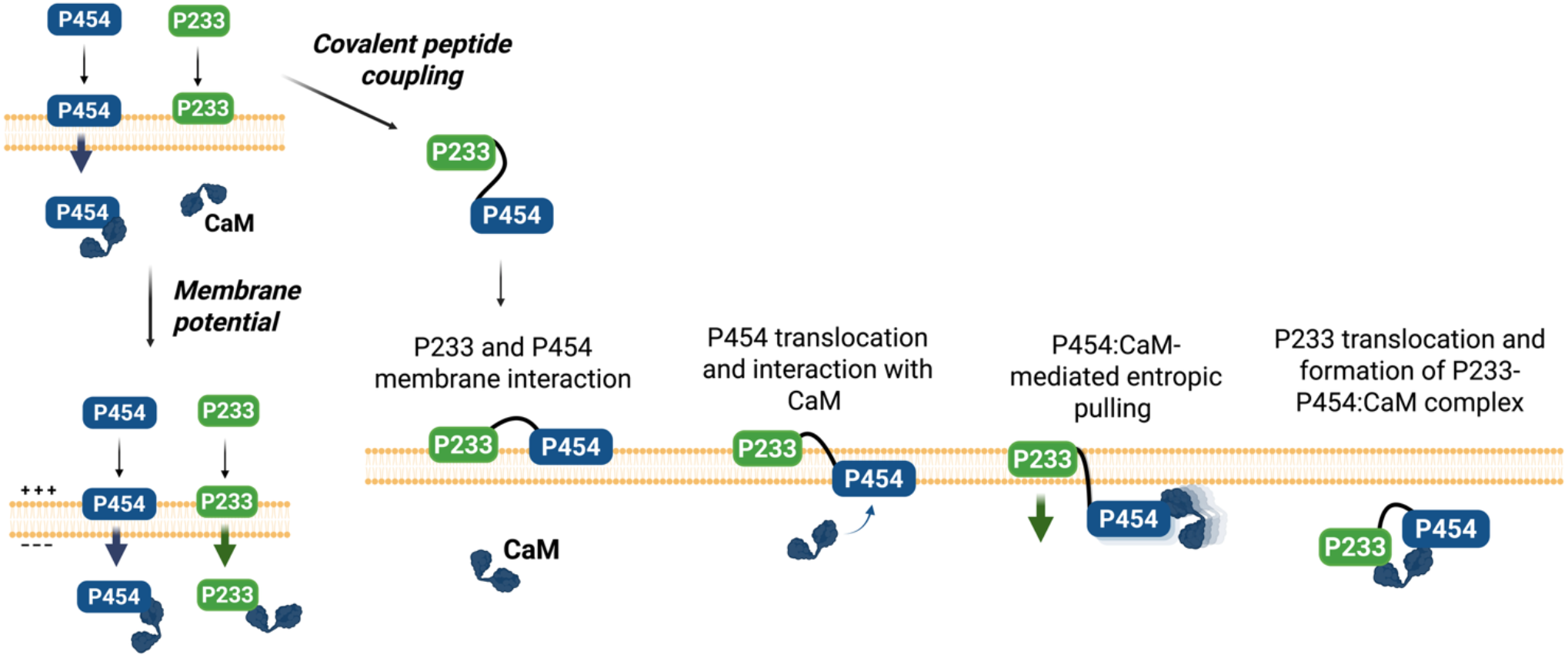
Schematic representation of the proposed mechanism of CyaA segments membrane translocation. In the presence of CaM in the *trans* compartment, the P454 peptide can translocate across membranes independently of membrane potential. In contrast, the P233 peptide behaves as a voltage-dependent membrane translocating segment strictly requiring a negative potential to translocate across the bilayer (left panel). A peptide containing both the P233 and P454 sequences is also capable of membrane translocation (right panel). We propose that, initially, both P454 and P233 segments interact with the membrane. P454 then translocates across the membrane, reaching the *trans* leaflet where it binds CaM. Formation of the CaM:P454 complex stabilizes the post-translocation state of P454 and generates an entropic pulling force transmitted along the polypeptide chain. This intramolecular force lowers the energy barrier for P233 translocation, enabling its translocation even in the absence of membrane potential. Upon translocation, P233 can also interact with CaM. Made with Biorender.

### Membrane potential tunes the translocation energy landscape of P233

The voltage dependence of P233 translocation suggests that membrane potential acts on P233 providing the energy needed to lower the effective free-energy barrier associated with membrane translocation. In this framework, membrane potential can actively modulate motion of positively charged peptides across membranes toward the *trans* side. The observation that P233 translocates at physiological membrane potential values underscores the relevance of this mechanism *in vivo* and is fully consistent with previous reports demonstrating the strict requirement of a negative membrane potential for ACD delivery into host cells^16,45^.

### Covalent linking enables coupled peptide translocation

A central finding of this study is that covalent coupling between P233 and P454 modulates P233 membrane translocation. While simple co-incubation of the two peptides does not promote P233 transport, the P233-P454 peptide, in which both sequences are connected by a nine-residue-long glycine-serine linker, readily translocates across membranes even in the absence of a membrane potential. From a biophysical perspective, this behavior indicates energetic coupling between the two segments: the spontaneous, CaM-dependent translocation of P454 effectively lowers the energetic barrier for the linked P233 peptide. This observation implies that the free energy released during P454 translocation and its subsequent extraction from the membrane by CaM can be transmitted along the polypeptide chain, generating a mechanical pulling force on P233. In this way, a local membrane-associated event is converted into a long-range force that drives the translocation of the covalently linked segment. This process is consistent with an entropic pulling^70–72^ mechanism that promotes vectorial transport of the polypeptide chain. This mode of operation is reminiscent of transport processes observed in protein translocation machineries^70–72^. Although no dedicated translocon is involved here, the high-affinity interaction between CaM and translocated P454 is sufficient to generate the entropic pulling force required to drive P233 translocation. The requirement for covalent linkage between P233 and P454 further demonstrates that this energy does not arise from simple co-localization of the two peptides at the membrane, but from direct traction of P233 by P454 through the peptide backbone.

### A coupled-force model for CyaA translocation

Taken together, our results support a mechanistic model in which CyaA translocation emerges from the concomitant action of multiple, partially independent driving forces (**Figure S3 - Supplementary Information)**. We propose that, following initial membrane binding and destabilization mediated by TR, HR and AR^32,33,73^, the P454 segment crosses the membrane, together with calcium^46^, engaging calcium-loaded CaM on the cytosolic side^47^. Binding of P454 to CaM is expected to selectively stabilize the post-translocation state, preventing back and forth transport of P454 across membrane. Yet, the experiments performed with the P233-P454 peptide presented here suggest that the spontaneous translocation of the P454 segments and its interaction with CaM can drive the translocation of its flanking regions by entropic pulling, thus lowering the energy barrier experienced by the N-terminal ACD. In parallel, the negative membrane potential may further decrease the energetic barrier for insertion and translocation of positively charged segments such as P233. The results presented here provide direct experimental support for a multicomponent translocation process : translocation arises from the coupling of peptide-membrane interaction, transmembrane negative electric potential, transmembrane calcium gradient and P454 cytosolic interaction with CaM. Once in the cytoplasm, ACD binding to CaM, primarily mediated by P233:CaM interactions^6^, triggers its folding and enzymatic activation^7,8^, which leads to cAMP overproduction and cellular intoxication^4,5^.

### The DIB-Pipette approach as a method to investigate force-driven membrane translocation

The DIB-Pipette represents a recently developed implementation of the DIB model membrane^59,60,62^ and is a useful approach to investigate membrane translocation events under precisely defined electrical conditions. Although previous works already successfully implemented membrane potential control in DIB experiments^61,63,74^, our approach combines the possibility of applying a transmembrane potential to a small droplet pair (droplet diameter ∼30 to 50 µm, *i*.*e*., volume ∼14 to 65 pL) with direct detection of membrane translocation by epifluorescence microscopy. Moreover, since the DIB-Pipette approach requires the formation of a single droplet pair each time, some major advantages in translocation detection are reported, compared to previously used methods^47,64^: the formation of a single, pipette-held droplet pair (*i*) minimizes background fluorescence and (*ii*) improves signal-to-noise ratio because the reduced dimension of droplets means a high membrane surface over droplet volume ratio. These two parameters enable reliable detection and high sensitivity in the measurement of rare translocation events. Therefore, by integrating the classical DIB readout of peptide translocation based on fluorescence detection^47,64^ with the reduced background fluorescence and improved signal-to-noise ratio induced by the single droplet pairs analysis and with direct voltage control, the DIB-Pipette approach represents a useful resource to perform membrane translocation studies.

## Conclusions

In conclusion, our results establish that P454 and CaM operate as an autonomous two-component membrane-translocating system, in which CaM binding converts P454 translocation into a directional process. In contrast, P233 behaves as a voltage-dependent segment whose translocation energy barrier is selectively lowered by a negative membrane potential (**Figure 7**). These two segments therefore operate under distinct energetic regimes: P454 intrinsically overcomes the membrane barrier, while P233 does not translocate unless an additional driving force is provided. A central finding of this work is that covalent coupling between the peptides enables P233 translocation even in the absence of membrane potential, demonstrating that membrane translocation can be driven by intramolecular force transmission (**Figure 7**). In this configuration, CaM binding to the translocated P454 segment stabilizes its post-translocation state and generates an entropic pulling force that is transmitted along the polypeptide chain, thereby lowering the free-energy barrier for P233 translocation.

More broadly, our findings suggest that CyaA membrane translocation is not encoded within a single region but instead emerges from the coordinated action of discrete, functionally specialized segments. In addition to established factors such as protein acylation, calcium gradient, and membrane potential, our results support a model in which CaM-dependent entropic pulling contributes directly to the initial steps of translocation by mechanically assisting the transport of CyaA membrane-active regions across the membrane. Finally, the DIB-Pipette provides a powerful and versatile approach to dissect how membrane potential, electrochemical gradients, membrane composition, peptide–membrane affinity, peptide physicochemical properties, and cytosolic binding partners are integrated to drive biomolecule translocation across membranes. The DIB-Pipette approach may also provide valuable insights into the translocation processes of various full-length toxins.

## Acknowledgements

This work was founded by Institut Pasteur (SPAIS/PTR PTR 2022-502, PTR 2023-708, PTR 2023-626 and DARRI/Emergence), CNRS and the Agence Nationale de la Recherche (3DTransCyaA project, ANR 21-CE11-0014). G.S. was supported by the Sorbonne University and Institut Pasteur PhD program. N.C. was supported by Institut Pasteur (DARRI/Emergence) and the ANR (ANR 21-CE11-0014). J.F. was funded by a Pasteur-Roux-Cantarini Tech fellowship (S-FB14002-85B) and the SPAIS/PTR 502-22 from Institut Pasteur. C.L. was supported by Institut Pasteur ANR (ANR 21-CE11-0014-01-3DTransCyaA). The 800-MHz NMR spectrometer of the Institut Pasteur was partially funded by the Région Ile de France (SESAME 2014 NMRCHR, grant 4014526). We acknowledge SOLEIL Synchrotron for provision of synchrotron radiation facilities.

