## Supplementary figures and table for "Covalently linked peptides and membrane potential enable CyaA segment translocation"

| | $K_d$ (µM) | $K_d$ error (µM) | $K_x$ | $\Delta G_{Kx}$ (kcal/mol) | $K_d$ (µM) | $K_d$ error (µM) | $K_x$ | $\Delta G_{Kx}$ (kcal/mol) |
|---|---|---|---|---|---|---|---|---|
| **Method** | Tryptophan fluorescence | Tryptophan fluorescence | Tryptophan fluorescence | Tryptophan fluorescence | B2LiVe | B2LiVe | B2LiVe | B2LiVe |
| **DOPC: POPG 8:2** | 582 | 46 | $9.53^{E}4$ | - 6.8 | 255 | 70 | $2.18^{E}5$ | - 7.3 |
| **POPC: POPG 8:2** | 1171 | 47 | $4.74^{E}4$ | - 6.4 | 462 | 94 | $1.20^{E}5$ | - 6.9 |
| **POPC** | 3095 | 378 | $1.79^{E}4$ | - 5.8 | 1127 | 804 | $4.92^{E}4$ | - 6.4 |

**Table S1:** Thermodynamic parameters of the solution to membrane partitioning of 5 µM of P233 obtained by tryptophan fluorescence and B2LiVe methods, at 25°C in the presence of increasing (0-10 mM) concentrations of DOPC:POPG 8:2 molar ratio, POPC:POPG 8:2 molar ratio and POPC 100% SUVs.

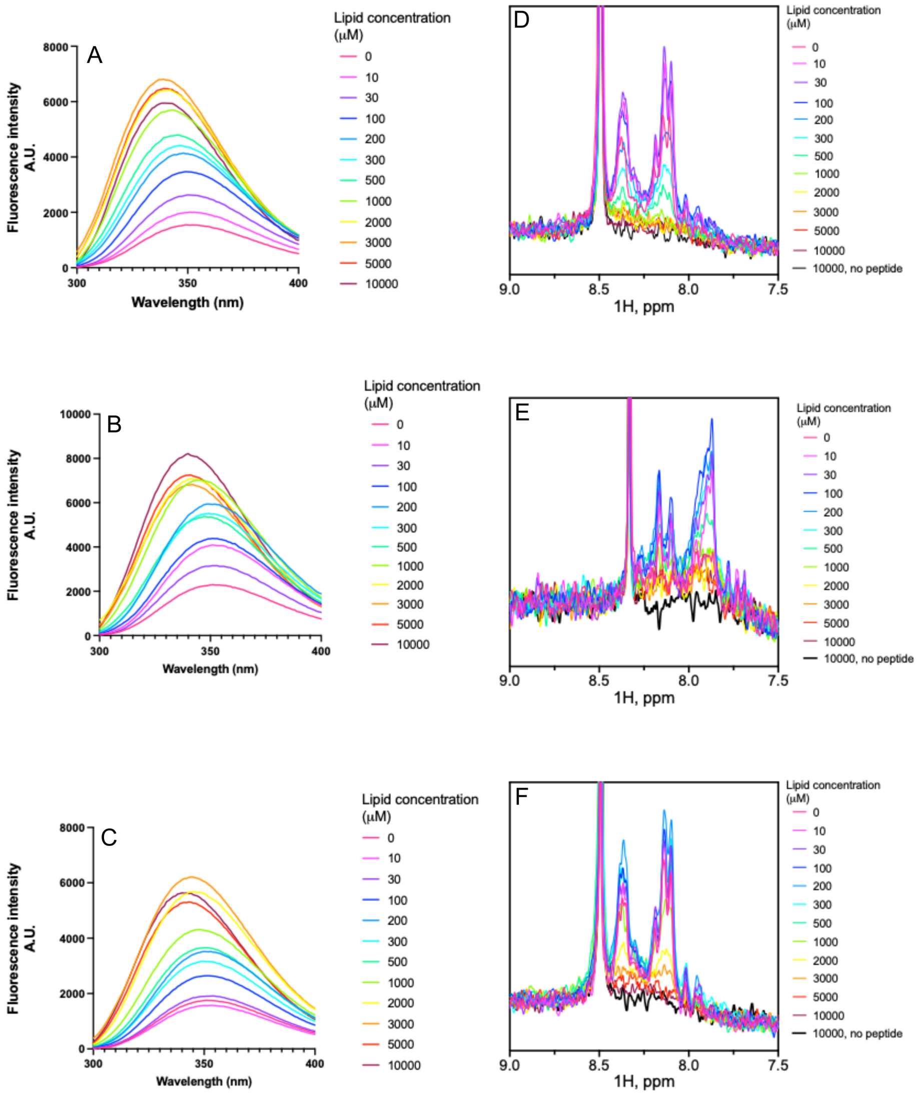


**Figure S1**: Solution to membrane partitioning of 5 µM of P233 monitored by changes of the tryptophan fluorescence (**A, B, C**), and by the B2LiVe method (**D, E, F**) at 25°C. The concentration of lipids ranged from 0 (in fuchsia) to 10 mM (in burgundy). Three different lipid compositions were tested: A) and D): DOPC:POPG 8:2, B) and E): POPC:POPG 8:2 and C) and F): POPC. In A, B, C membrane partitioning is indicated by a shift towards lower wavelength values of the maximum fluorescence emission wavelength and by an increase of the fluorescence intensity. In D, E, F membrane partitioning is followed by disappearance of the amide resonance signal. The sharp peak around 8.5 ppm is due to a trace impurity of formic acid contained in $D_2O$. The spectrum of SUVs in the absence of peptide is reported as a black line.

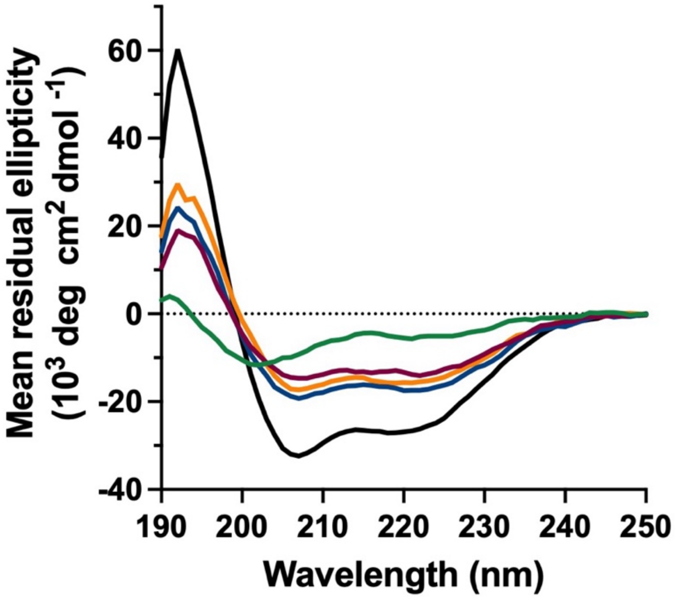


**Figure S2:** Far UV Synchrotron radiation circular dichroism (SRCD) spectra of 100 µM P454 and 100 µM P233 recorded in the absence (green line) and in the presence of 6 mM SUVs composed of POPC (burgundy line), POPC:POPG at an 8:2 molar ratio (blue line), or DOPC:POPG at an 8:2 molar ratio (orange line), as well as in the presence of 40% TFE (black line) in buffer A at 25 °C.

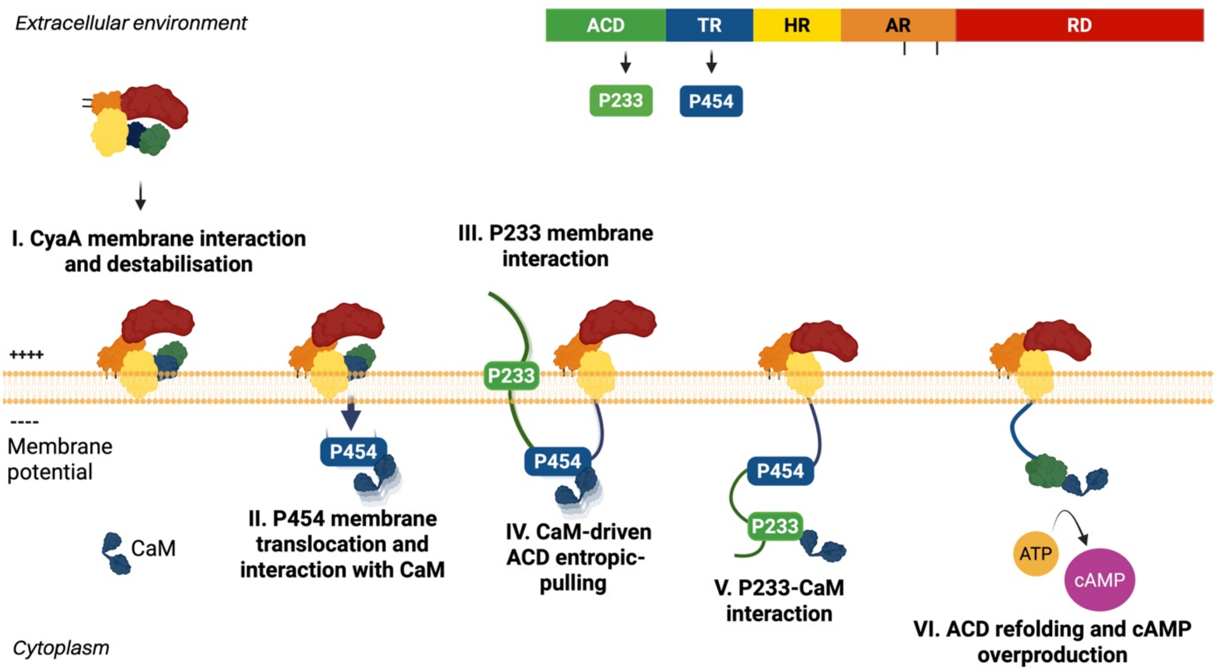


**Figure S3:** Schematic representation of the proposed mechanism of CyaA membrane translocation. Initially, CyaA interacts with the target cell membranes and induces membrane destabilization (I). The P454 region interacts with the plasma membrane and, independently of the presence of a negative membrane potential, translocates across the lipid bilayer. Once in the *trans* compartment of the membrane, P454 binds to CaM, that pulls TR inside the cell cytoplasm (II). The P233 segment, upon membrane interaction docks its flanking regions from ACD to the membrane (III). P454 translocation may facilitate the translocation of other protein segments: we propose that formation of the P454:CaM complex generates an entropic pulling force that promotes ACD unfolding and drives its translocation (IV). Supported by membrane potential and by the entropic pulling force produced by the P454:CaM complex, the P233 region translocates and binds to CaM (V). ACD achieves its translocation. ACD folds upon CaM binding and overproduces cAMP from cellular ATP at high turnover rate (VI). CyaA organization is reported on the upper left section of the figure. CyaA domains are represented as colored rectangles: RD in red, AR in orange, HR in yellow, TR in blue, ACD in green. P454 is represented as a blue rectangle, P233 is represented as a green rectangle. CaM is shown in dark blue. The membrane is represented in yellow. ATP is represented as a dark yellow circle, cAMP as pink circle. Molecules are not at scale. Made with Biorender
